# Reconciling disparate estimates of viral genetic diversity during human influenza infections

**DOI:** 10.1101/364430

**Authors:** Katherine S. Xue, Jesse D. Bloom

## Abstract

Deep sequencing can measure viral genetic diversity within human influenza infections, but published studies disagree in their estimates of how much genetic diversity is typically present. One large-scale deep-sequencing study of human influenza reported high levels of shared viral genetic diversity among infected individuals in Hong Kong, but subsequent studies of other cohorts have reported little shared viral diversity. We re-analyze sequencing data from four studies of within-host genetic diversity encompassing more than 500 acute human influenza infections. We identify an anomaly in the Hong Kong data that provides a technical explanation for these discrepancies: read pairs from this study are often split between different biological samples, indicating that some reads are incorrectly assigned. These technical abnormalities explain the high levels of within-host variation and loose transmission bottlenecks reported by this study. Studies without these anomalies consistently report low levels of genetic diversity in acute human influenza infections.

## Main text

A key question in the study of influenza-virus evolution is how rapidly viral genetic variation arises within infected humans, and how much of this genetic diversity is maintained during transmission (McCrone and Lauring, 2018; Xue et al., 2018a). Recently, several studies have measured influenza’s within-host genetic diversity in large cohorts of infected humans using high-throughput deep sequencing (**Supplemental Table 1**) (Debbink et al., 2017; Dinis et al., 2016; McCrone et al., 2018; Poon et al., 2016). These studies have disagreed considerably in their estimates of influenza’s within-host genetic diversity.

One of the first large-scale studies of influenza’s genetic diversity in human infections analyzed a household cohort in Hong Kong (Poon et al., 2016). This study reported high within-host genetic diversity, estimating that approximately 40% to 66% of patients harbor mixed infections, resulting from co-infection with multiple genetically distinct viral lineages. However, subsequent deep sequencing studies of influenza’s genetic diversity in human cohorts from Wisconsin (Dinis et al., 2016), Michigan (Debbink et al., 2017; McCrone et al., 2018), and Washington (Xue et al., 2018b) have reported lower levels of viral genetic diversity, with few mixed infections. One important reason to measure within-host viral diversity is to estimate transmission bottleneck sizes (McCrone and Lauring, 2018; Sobel Leonard et al., 2017; Xue et al., 2018a), and the wide variation in estimates of viral genetic diversity has led to wide variation in estimates of bottleneck sizes. The Hong Kong study estimated a bottleneck of 200–250 viral genomes (Poon et al., 2016; Sobel Leonard et al., 2017), while a subsequent Michigan study estimated a bottleneck of just to 1–2 viral genomes (McCrone et al., 2018). These large discrepancies between major published studies are concerning.

Various biological and technical explanations could account for these discrepancies. The studies differ in their geographic locations, cohort design, deep sequencing methods, and data analysis approaches. Any of these factors could affect estimates of viral genetic diversity (Djikeng et al., 2008; Illingworth et al., 2017; McCrone and Lauring, 2016; Sobel Leonard et al., 2017). Here, we examine whether technical differences in the underlying deep-sequencing datasets or the methods used to analyze them explain the disparate estimates of within-host viral genetic diversity.

To systematically compare the results across studies, we used the same computational framework to re-analyze raw sequencing data for four large-scale studies of influenza’s within-host genetic diversity, together encompassing more than 500 acute human infections (Debbink et al., 2017; Dinis et al., 2016; McCrone et al., 2018; Poon et al., 2016). For each study, we applied the same variant-calling thresholds as the Hong Kong study, identifying sites with a minimum coverage of 200 at which a non-consensus base exceeds a frequency of 3% in the sequenced reads at that site (**Materials and methods**). We averaged variant frequencies between sequencing replicates where available but otherwise used an analysis pipeline that was as similar as possible across studies to ensure comparable estimates of within-host genetic diversity.

Our analysis recapitulates the major results reported in the Hong Kong study. **Supplemental Figure 1** shows within-host variation in the hemagglutinin gene in H3N2 patients in our re-analysis of the study’s data, in the same format as the second figure of the original publication (Poon et al., 2016). In both the original study and our re-analysis, the same within-host variant is often present at similar frequencies in multiple, epidemiologically unrelated individuals. Moreover, the minority variant in one group of samples is typically the majority or consensus variant in the remaining samples (**Supplemental Figure 1A**). Across the hemagglutinin gene, the original Hong Kong study and our re-analysis of that study’s data identify the same patterns of within-host variation (**Supplemental Figure 1B**).

Our analysis also identifies major differences between the Hong Kong dataset and the other studies. We find little within-host viral variation in the other three datasets, in line with these studies’ stated conclusions (**Figure 1A**) (Debbink et al., 2017; Dinis et al., 2016; McCrone et al., 2018). Furthermore, only the Hong Kong dataset contains high-frequency within-host variants that are shared between epidemiologically unrelated individuals. In data from the Hong Kong study, the same within-host variants were shared among more than half of the patients at 42 sites in the H3N2 genome, and 9 sites in the pdmH1N1 genome (**Figure 1B**). In contrast, we identified no such sites of extensively shared genetic variation among patients in the other three studies. These results show that the large discrepancies between the Hong Kong study and other published work cannot be accounted for solely by methodological differences in variant calling pipelines.

**Figure 1.**
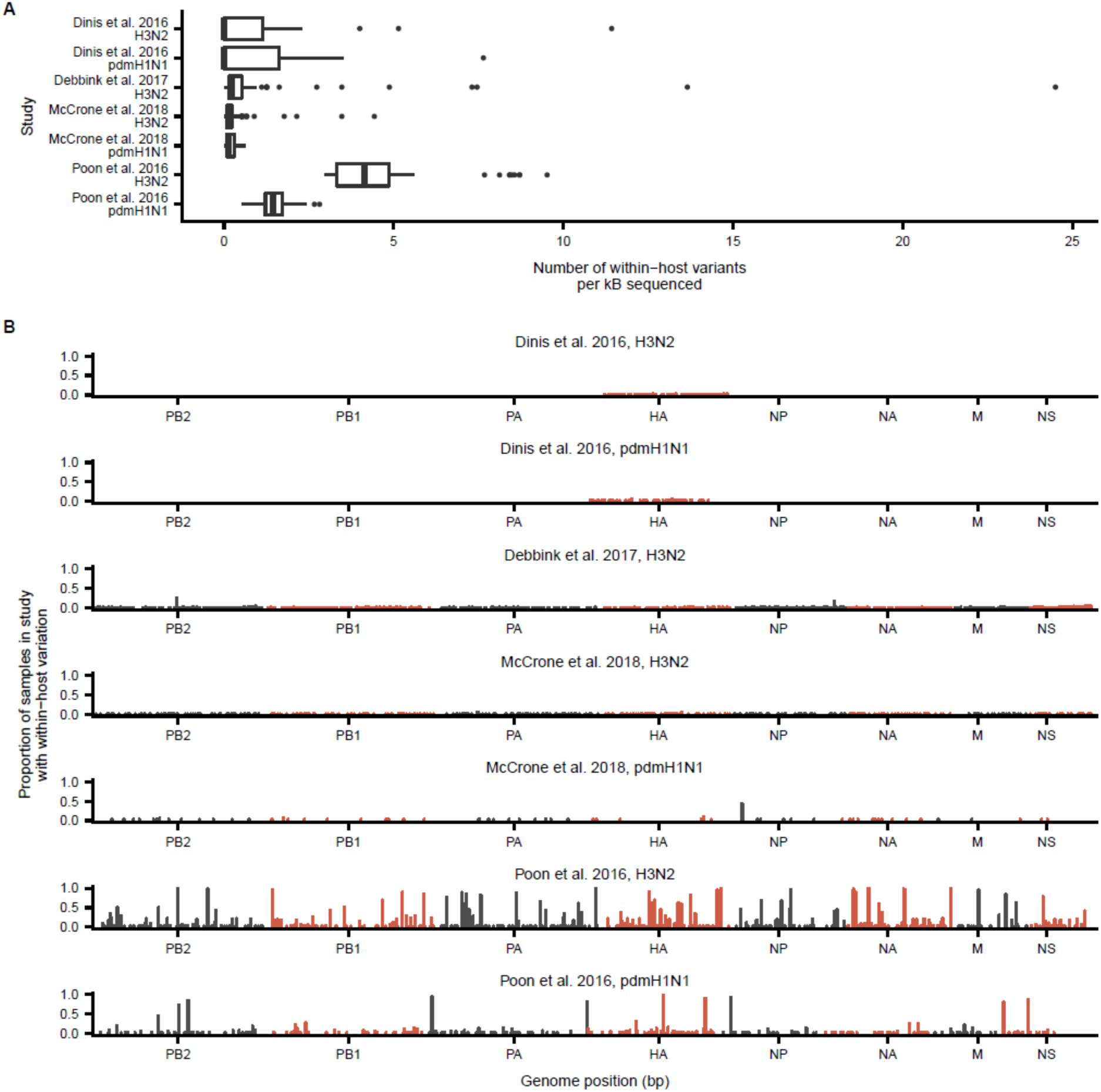
Comparison of within-host viral genetic diversity in four large-scale deep-sequencing studies of human influenza virus. **(A)** Number of within-host variants identified in each sample in each study, normalized to the length of the genome sequenced in each study. For each sample, we identified within-host variants that were present at a frequency of at least 3% at sites with minimum sequencing coverage of 200 reads. **(B)** Proportion of samples in each study in which we identified within-host variation at each genome site. Our re-analysis is consistent with the previously reported results of each study: we find little shared genetic diversity in the data from the Dinis et al. (2016), Debbink et al. (2017), and McCrone et al. (2018) studies, but we observe high shared genetic diversity in the data from the Poon et al study.

The extensive shared genetic diversity in the Hong Kong study could result from genuine similarity in the mix of viruses that infect epidemiologically unrelated humans in Hong Kong. But they could also reflect cross-contamination or other abnormalities in the underlying sequencing data. In the course of our analysis, we identified abnormalities in the raw sequencing data from the Hong Kong study that can explain the apparently high levels of shared viral genetic diversity across different infected individuals. The deep sequencing for this study used paired-end Illumina reads. Both reads in a pair come from the same molecule of PCR-amplified viral genetic material, and so should always be assigned to the same infected human (**Figure 2A**). Illumina software assigns standard headers to each FASTQ-format sequencing read. These header lines contain information about each read, including the sequencing lane, a unique read-pair identifier, and whether a read is the first or second member of a pair (**Figure 2B**). When we analyzed FASTQ headers in the raw sequencing data for the Hong Kong study, we found that paired-end sequencing reads were frequently split between samples assigned to different individuals (**Figure 2C**). (**Figure 1** and **Supplemental Figure 1** were generated by analyzing the sequencing data from the Hong Kong study as single-end data.) For instance, the read @SOLEXA4_0078:1:1101:10000:101622#ATCACG/1 was associated with study subject 737-V1 (0), whereas its pair @SOLEXA4_0078:1:1101:10000:101622#ATCACG/2 was associated with study subject 741-V1(0), an epidemiologically unrelated individual.

**Figure 2.**
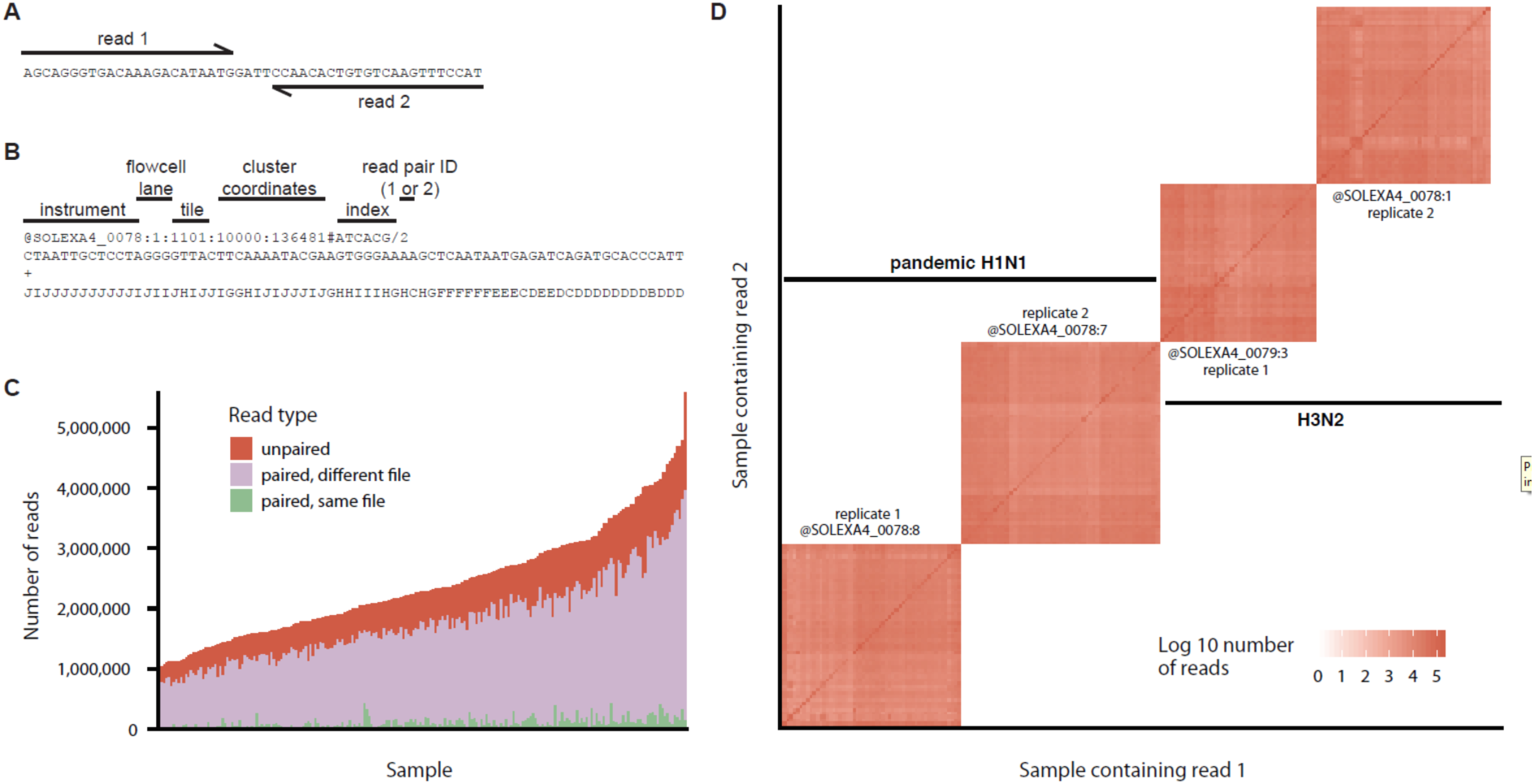
Paired-end sequencing reads are frequently split between samples that were run on the same sequencing lane. **(A)** Paired-end sequencing reads are derived from the same physical DNA molecule. **(B)** The FASTQ header for each sequencing read provides information about the sequencing instrument, flowcell lane, tile, cluster coordinates, and sequencing index for each read, as well as whether the read is the first or second member of a read pair. **(C)** Sequencing reads from the Hong Kong dataset are frequently split between distinct biological samples. **(D)** Hierarchical clustering of the number of read pairs split between each pair of samples in the Hong Kong study. Sequencing reads from the Hong Kong dataset are split between four distinct clusters of samples. All sequencing reads in each cluster are derived from the same flowcell lane and correspond to one set of replicate samples for one of the two influenza subtypes sequenced in the study.

It is biologically impossible for reads in a pair to be associated with distinct individuals, since both reads originate from the same DNA molecule. Across all samples, 70% of reads had corresponding pairs in a FASTQ file assigned to a different individual, and 25% of reads were not part of an identifiable pair (**Figure 2C**). Only 5% of the 500 million sequencing reads in this study were associated with the same sample as their corresponding pairs. This splitting of read pairs between samples indicates a problem in the sample index de-multiplexing or downstream computational analysis, and can be considered a form of technical cross-contamination.

Importantly, the problem appears to be with how read pairs were assigned to samples rather than with the FASTQ headers. We found that 93% of the read pairs reconstructed based on FASTQ header information mapped concordantly to the H3N2 or pandemic H1N1 influenza genome—that is, both reads in a pair mapped to the same gene segment in the expected relative orientation.

We analyzed patterns of read-pair splitting between all samples in the study (**Figure 2D**). We identified four disjoint sets of samples for which read pairs are split extensively within sets, but never between sets. Further analysis of FASTQ headers showed that all of the sequencing reads from each cluster were derived from the same flowcell lane. Poon et al. 2016 report that samples were amplified in duplicate and that replicates were sequenced on distinct flowcell lanes. Indeed, we find that each set of samples corresponds almost exactly to one set of replicate samples for one of the two influenza subtypes sequenced in this study (**Figure 2D**). This finding was robust to the computational analysis pipeline: the first author generated all of the figures in this paper (https://github.com/ksxue/compare-flu-within-hosts-public), but the last author conducted an independent re-analysis of the data to reach similar conclusions (https://github.com/jbloomlab/reanalyze_Poon_et_al). Altogether, these analyses suggest that read pairs are split extensively between samples of a given influenza subtype in the Hong Kong study.

Without access to the full computational pipeline for the Hong Kong study, we cannot determine directly whether the first read, second read, or both members of split read pairs were assigned to samples incorrectly. However, when we analyzed only the first read of each pair, we found low within-host diversity, in line with other studies (**Figure 3**). In contrast, the second read of each pair was responsible for the high viral diversity reported in the Hong Kong study. These results suggest that the second member of each read pair may have been incorrectly assigned, and the first member may more accurately represent the low levels of within-host viral diversity.

**Figure 3.**
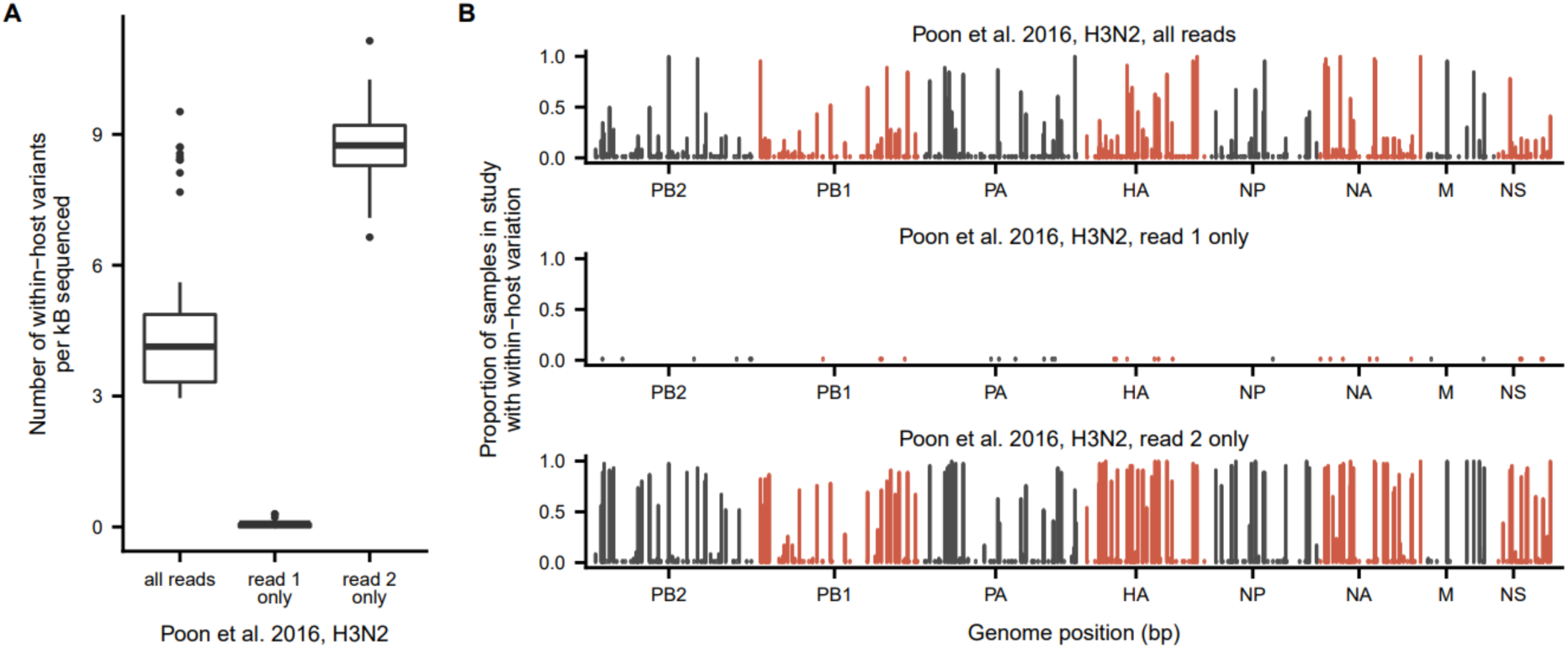
Within-host genetic diversity in the Hong Kong study is primarily located on read 2. **(A)** Number of within-host variants identified in the H3N2 samples when analyzing both members of each read pair, just read 1, or just read 2. For each sample, we identified within-host variants that were present at a frequency of at least 3% at sites with minimum sequencing coverage of 200 reads. **(B)** Proportion of samples for which we identified within-host variation at each genome site when analyzing both reads for a pair, just read 1, or just read 2.

This splitting of read pairs between unrelated samples has important consequences for estimates of viral genetic diversity within human infections. Even if each individual were infected with a clonal population of influenza virus, read-pair splitting would create the appearance of high levels of shared genetic diversity between unrelated individuals. For instance, at a site in the influenza genome where some individuals exclusively have nucleotide A and others exclusively have nucleotide T, read-pair splitting would make it seem as though all individuals with majority identity A have minority variant T and vice versa, even in the absence of genuine within-host variation. The high-frequency shared viral diversity within human hosts in the Hong Kong study corresponds closely to what would be expected from read-pair splitting (**Supplemental Figure 1A**), suggesting that this abnormality may be responsible for the published results.

Read-pair splitting may also explain why the Hong Kong household cohort study estimates a loose transmission bottleneck for human influenza virus of 200–250 viral genomes (Poon et al., 2016; Sobel Leonard et al., 2017), compared to a Michigan household cohort study that estimates a bottleneck size of 1–2 viral genomes (McCrone et al., 2018). Splitting of read pairs between samples would also create the appearance of shared within-host variation in donor and recipient individuals in a transmission chain, resulting in estimates of a looser transmission bottleneck.

Our finding of read-pair splitting in the Hong Kong dataset provides a technical explanation for major discrepancies in recent studies of the genetic diversity of human influenza viruses. In particular, these technical anomalies may account for the Hong Kong study’s finding of shared, high-frequency viral diversity within human hosts (Poon et al., 2016) and its estimate of a loose transmission bottleneck between hosts (Poon et al., 2016; Sobel Leonard et al., 2017). If we exclude the Hong Kong study, then all other studies report low levels of within-host genetic diversity for human influenza virus (Debbink et al., 2017; Dinis et al., 2016; McCrone et al., 2018).

## Materials and methods

### Code availability

The computer code that performs the analysis is available at https://github.com/ksxue/compare-flu-within-hosts-public. All figures were generated by the first author using this code base, but the last author independently conducted an analysis of read pairing and came to similar conclusions (https://github.com/jbloomlab/reanalyze_Poon_et_al).

### Source data

We downloaded sequencing data generated by the Hong Kong study (Poon et al., 2016) from https://www.synapse.org/#!Synapse:syn8033988, following the methods of a study that re-analyzed data from the Hong Kong study to estimate transmission bottleneck sizes using a new analytical method (Sobel Leonard et al., 2017). We obtained sequencing data for the Wisconsin study (Dinis et al., 2016) by personal communication. We downloaded sequencing data for the other studies from SRA BioProject PRJNA344659 (Debbink et al., 2017) and PRJNA412631 (McCrone et al., 2018).

### Variant calling

Here, we summarize our general computational pipeline for variant calling, with modifications for individual studies described below. We used cutadapt 1.8.3 (Martin, 2011) to trim Nextera adapter sequences, remove bases at the ends of reads with a Q-score below 24, and filter out reads whose remaining length was shorter than 20 bases. To determine the sample subtype, we used bowtie2 to map 1000 reads from each sample to reference genomes for each influenza subtype: A/Victoria/361/2011 (H3N2), A/California/04/2009 (pdmH1N1), and A/Boston/12/2007 (seasonal H1N1). We determined what proportion of the reads from each sample mapped to each reference genome and classified sample subtype based which reference genome resulted in the highest mapping rate.

For each sample, we first aligned reads to the subtype reference genome using bowtie2 and the --`very-sensitive` setting (Langmead and Salzberg, 2012), then we used custom scripts to tally the counts of each base at each genome position and infer a consensus sequence for that sample. We then realigned the reads to the sample consensus sequence, discarding reads with a mapping score below 20 and bases with a Q-score below 20, and we removed read duplicates using Picard version 1.43. We used custom scripts to tally the counts of each base at each genome position among the remaining reads.

We defined sites of within-host variation as positions in the genome with sequencing coverage of at least 200 reads at which a minority base is present at a frequency of at least 3%, following the variant-calling criteria of the Hong Kong study (Poon et al., 2016). The Hong Kong study sequenced most samples in duplicate, and we required within-host variants to be present at a frequency of 3% in both sequencing replicates. To maintain consistent variant-calling criteria for all samples in the Hong Kong study, we did not include samples with only a single sequencing replicate in our downstream analyses. We performed no other filtering for sample or variant quality because common filtering metrics like sequencing replicates (Xue et al., 2017) and plasmid sequencing controls (McCrone and Lauring, 2016; McCrone et al., 2018) were not universally available.

### Study-specific modifications

We tried to analyze all sequencing data using methods that were as similar as possible across studies. However, certain study designs or data formats required us to modify our basic analysis framework as described below. For more information, the code that performs the analysis is available at https://github.com/ksxue/compare-flu-within-hosts-public.

#### Hong Kong study

We obtained data from the Hong Kong study (Poon et al., 2016) from https://www.synapse.org/#!Synapse:syn8033988 in the form of a single FASTQ file per biological sample containing first and second members of read pairs. As described in the main text and in the section below, we found that read pairs were frequently split between different sample files, so we could not conduct a meaningful analysis of paired-end sequencing reads. To call variants and generate the data in Figures 1 and 2, we analyzed the sequencing data as single-end reads. We used single-end read-mapping settings for bowtie2, and we did not perform read deduplication, which typically makes use of paired-end read information. Most samples from this study were sequenced in duplicate. We only analyzed samples for which we were able to identify both sequencing replicates, and we required within-host variants to meet the variant-calling criteria in both replicates.

#### Wisconsin study

We obtained data from the Wisconsin study (Dinis et al., 2016) by personal communication in the form of a single FASTQ file per biological sample containing first and second members of read pairs. We reconstructed read pairs for each sample using read-pair information in the FASTQ headers, and we found no read pairs that were split between different sample files. We then analyzed the reconstructed read pairs as described above.

### Analysis of read pairing

To analyze read pairing in data from the Hong Kong study, we parsed FASTQ read headers to identify first and second members of read pairs, as well as their associated sequencing lane. We interpreted the colon-delimited fields of a FASTQ header like @SOLEXA4_0078:1:1101:10000:101622#ATCACG/1 to contain information about the sequencing instrument (@SOLEXA4_0078), lane number (`1`), tile (`1101`), cluster coordinate (`10000:10162 2`), sequencing index (`ATCACG`), and whether a read was the first or second member of a read pair (`1`), in accordance with Illumina specifications (for instance, see http://support.illumina.com/content/dam/illumina-support/help/BaseSpaceHelp_v2/Content/Vault/Informatics/Sequencing_Analysis/BS/swSEQ_mBS_FASTQFiles.htm). We did not make use of sequencing indices in this analysis. We used this FASTQ information to analyze all 500 million reads in the Hong Kong dataset to determine what proportion of reads had pairs in the same sample file and to determine which reads were associated with each sequencing lane. We used the `--very-fast-local` setting of bowtie2 to map reconstructed read pairs against a pan-influenza reference genome produced by concatenating the eight-segment A/Victoria/361/2011 (H3N2) reference genome and the eight-segment A/California/04/2009 (pdmH1N1) reference genome into a single reference genome containing sixteen segments.

### Analysis of first and second reads

To separate the first and second reads assigned to each sample file in the Hong Kong dataset, we interpreted the colon-delimited fields of the FASTQ headers as described above. Because read pairs were frequently split between sample files, the first and second reads that we identified in each sample did not necessarily constitute complete pairs. We then used the read-mapping and variant-calling pipeline described above for the Hong Kong study to identify sites of within-host variation.

## Acknowledgments

We thank Phil Green for helpful comments on the manuscript. KSX is supported by the Hertz Foundation Myhrvold Family Fellowship. The work of JDB was supported by grant R01AI127893 from the NIAID of the NIH and a Faculty Scholars grant from HHMI and the Simons Foundation.

## SUPPLEMENTAL FIGURES

**Supplemental Figure 1.**
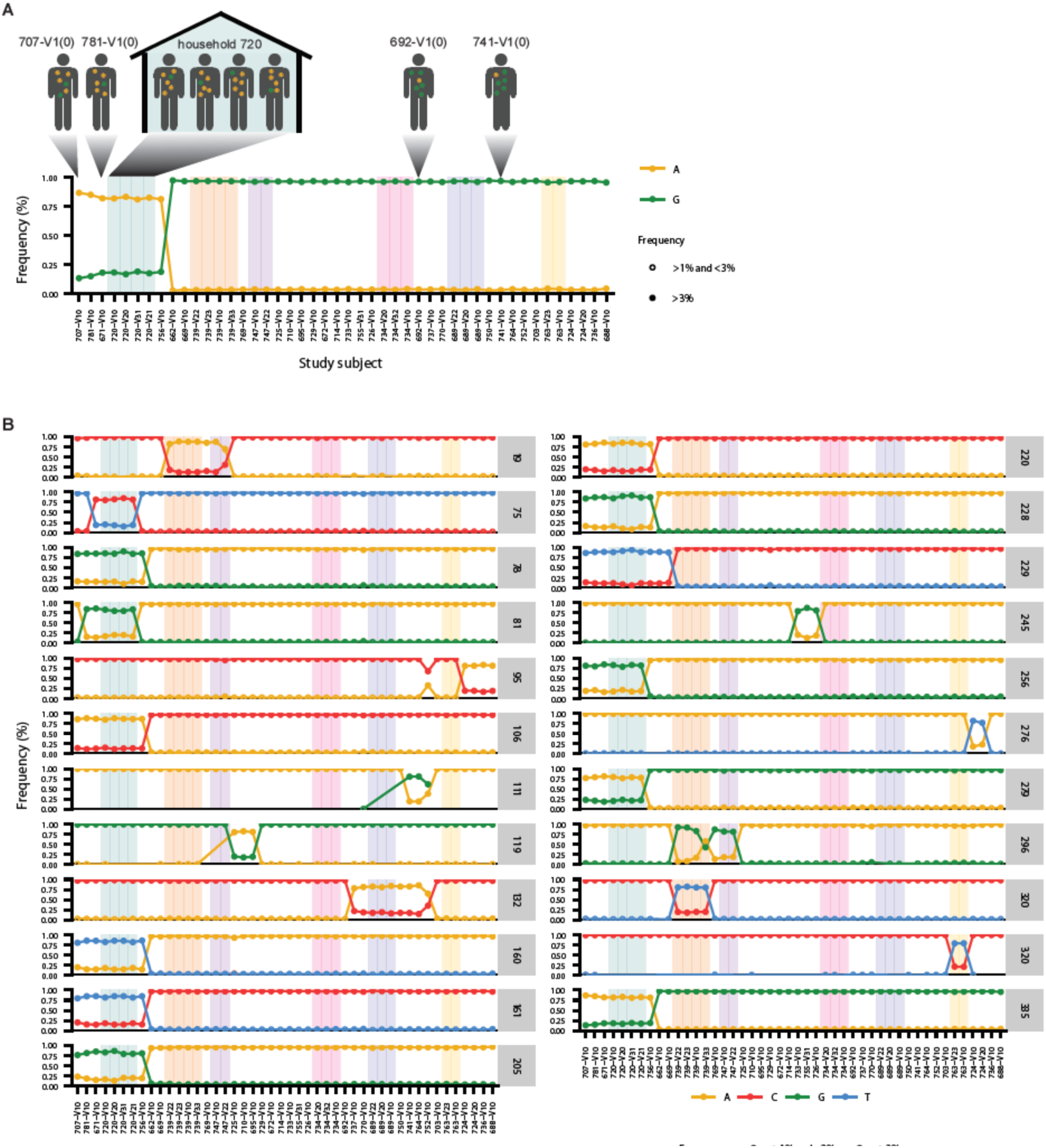
High within-host genetic diversity of human influenza virus in our re-analysis of sequencing data from the Hong Kong study. This figure mimics the format of the second figure of Poon et al (2016) and shows that our re-analysis recapitulates the main reported results of high-frequency shared genetic diversity between epidemiologically unrelated individuals. **(A)** Viral genetic diversity at hemagglutinin codon 335 in H3N2 human influenza infections. At this codon, both variants encode the same amino acid. This plot shows within-host variants that were present at a frequency of at least 1% in both sequencing replicates at sites with minimum sequencing coverage of 200 reads. Shaded regions indicate individuals from the same household. **(B)** Viral genetic diversity in the HA1 domain of hemagglutinin in H3N2 human influenza infections in our re-analysis. Each panel represents a separate site in the genome and is labeled by the HA codon it represents. Sites shown harbored within-host variation at a frequency of at least 3% in both sequencing replicates for at least two samples in the study.

**Supplemental Table 1.**
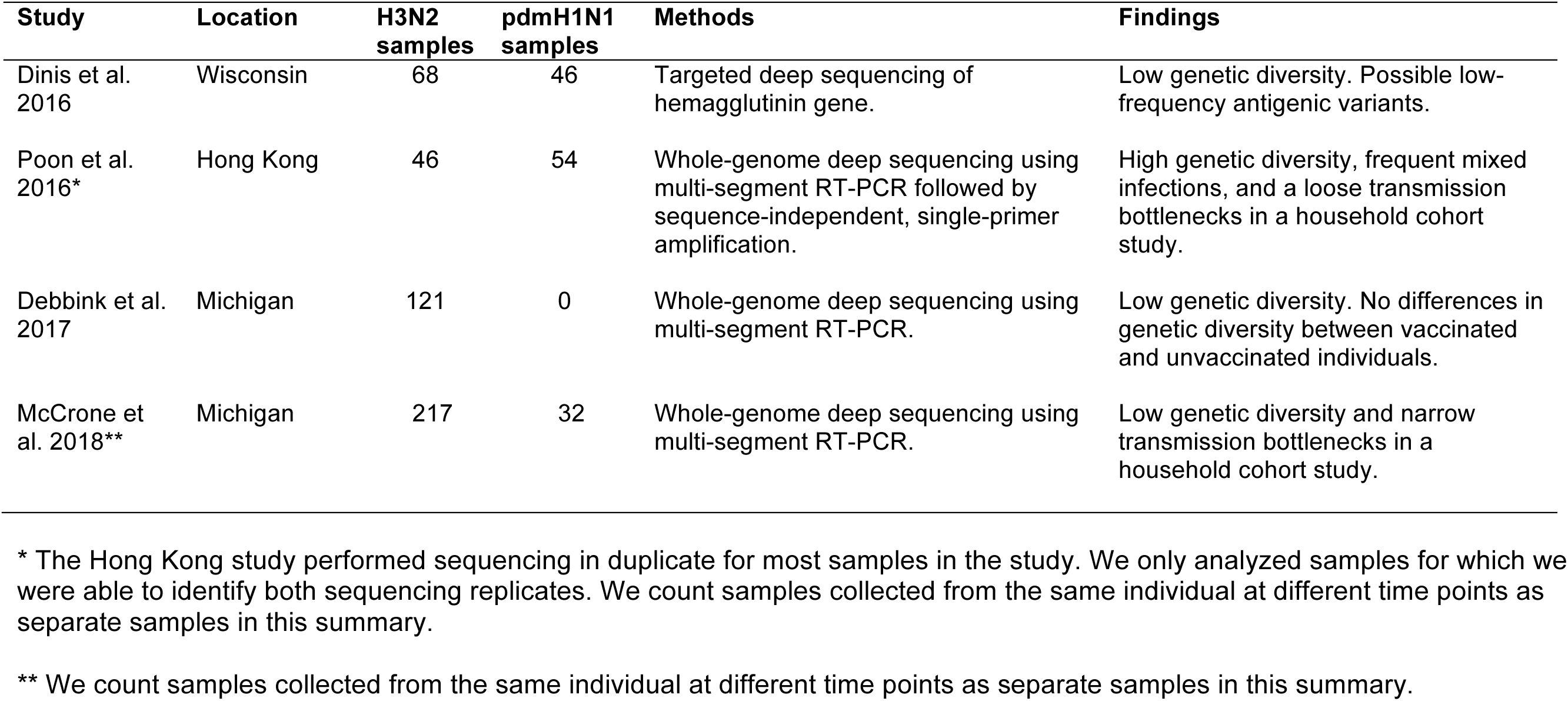
Large-scale deep-sequencing studies of human influenza virus.

